# A complete and dynamic tree of birds

**DOI:** 10.1101/2024.05.20.595017

**Authors:** Emily Jane McTavish, Jeff A. Gerbracht, Mark T. Holder, Marshall J. Iliff, Denis Lepage, Pam Rasmussen, Benjamin Redelings, Luna Luisa Sanchez Reyes, Eliot T. Miller

## Abstract

We present a complete, time-scaled, evolutionary tree of the world’s bird species. This tree unites phylogenetic estimates for 9,239 species from 262 studies published between 1990 and 2024, using the Open Tree synthesis algorithm. The remaining species are placed in the tree based on curated taxonomic information. The tips of this complete tree are aligned to the species in the Clements Taxonomy used by eBird and other resources, and cross-mapped to other taxonomic systems including the Open Tree of Life (Open Tree), National Center for Biotechnology Information (NCBI), and Global Biodiversity Information Facility (GBIF). The total number of named bird species varies between 10,824 and 11,017 across the taxonomy versions we applied (v2021, v2022 and v2023). We share complete trees for each taxonomy version. The procedure, software and data-stores we used to generate this tree are public and reproducible. The tree presented here is Aves v1.2 and can be easily updated with new phylogenetic information as new estimates are published. We demonstrate the types of large scale analyses this data resource enables by linking geographic data with the phylogeny to calculate the regional phylogenetic diversity of birds across the world. We will release updated versions of the phylogenetic synthesis and taxonomic translation tables annually. The procedure we describe here can be applied to developing complete phylogenetic estimates for any taxonomic group of interest.

**Significance statement:** Birds are charismatic - well loved, and highly studied. Many new phylogenies elucidating avian birds evolutionary relationships are published every year. We have united phylogenetic estimates from hundreds of studies to create a complete evolutionary tree of all birds. While a variety of resources aggregate huge collections of trait, behavior and location data for birds, previously the barriers to linking data between these data resources and bird evolutionary history have limited the opportunities to do exciting large scale analyses. We have bridged that gap, and developed a system that allows us to easily update our understanding of bird evolution as new estimates are generated. We share a workflow and the software needed to create a complete evolutionary tree for any group.

## Introduction

Advances in data availability make it increasingly possible to address large scale biological questions across the diversity of life. More researchers and project teams are providing open access to their primary data, which has accelerated the progress of integrative research. However, to build on this momentum, it is necessary to not only make data available, but also to make it accessible and dynamic. The tree of life provides a comprehensive, biologically relevant means to link data sets, and cohesively analyze data from multiple sources. Historically, the lack of a unified framework for accessing evolutionary trees has resulted in a gap between the novel inferences of relationships between species generated by systematists, and the analyses researchers want to perform using the tree of life [7, 19]. Large scale estimates of biodiversity have relied on outdated taxonomic relationships, rather than cutting edge evolutionary inferences [13].

The Open Tree of Life project (Open Tree, hereafter) bridges that gap by providing phylogenetically informed estimates of relationships for taxa across the entire tree of life [10, 19]. Open Tree provides a framework for linking names, metadata, and phylogenetic information across the entire tree of life. Since the publication of the first draft tree in 2015 [10], Open Tree has developed into a robust biodiversity informatics resource. Here, we apply the synthetic tree building and taxonomy tools developed by Open Tree to provide a complete and updatable, time-scaled, tree of all bird species.

This project not only provides a complete, phylogenetically rigorous tree of all bird species, it also provides a framework for maintaining an accurate and readily available estimate of bird relationships, even as new species are discovered, and prior relationships and taxonomy are revised. The input data curation and the published data products all follow FAIR principles for data stewardship - Findability, Accessibility, Interoperability, and Reuse of digital assets [35]. All the input phylogenies are available and have extensive curated metadata [20]. The phylogenies in the data store are searchable by their publication information as well as indexed by the phylogenetic content itself, using taxonomic context. We developed and published taxonomic translation tables which make this tree estimate easily connected with large existing bird data sets, including AVONET [32] trait data and eBird distribution data [29], among many others. This tree incorporates the relationships published in 281 bird phylogenies published in 262 studies from 1990 to 2024.

All data and code are publicly available, and the procedure to generate the tree is fully reproducible. The tree and taxonomic translation tables are versioned, and will be updated annually. This synthesis framework can also be applied to developing and maintaining a complete dynamic phylogenetic estimate for any taxonomic group of interest.

## Results

We unified 281 input phylogenies into a phylogenetic synthesis for 9,239 species across all birds (Figure 1, 2). We used curated taxonomic information to add in taxa which were absent from phylogenetic estimates to generate complete trees with all species present in the 2021, 2022, and 2023 Clements taxonomy versions (Figure 1, Table 1). The provenance of each branch can be traced back to the input studies which support that relationship (Figure 2a). The tip labels on these species-level trees can be easily translated between Open Taxonomy, NCBI, GBIF and Clements taxonomy labels using the unique identifiers associated with each taxonomy.

**Figure 1.**
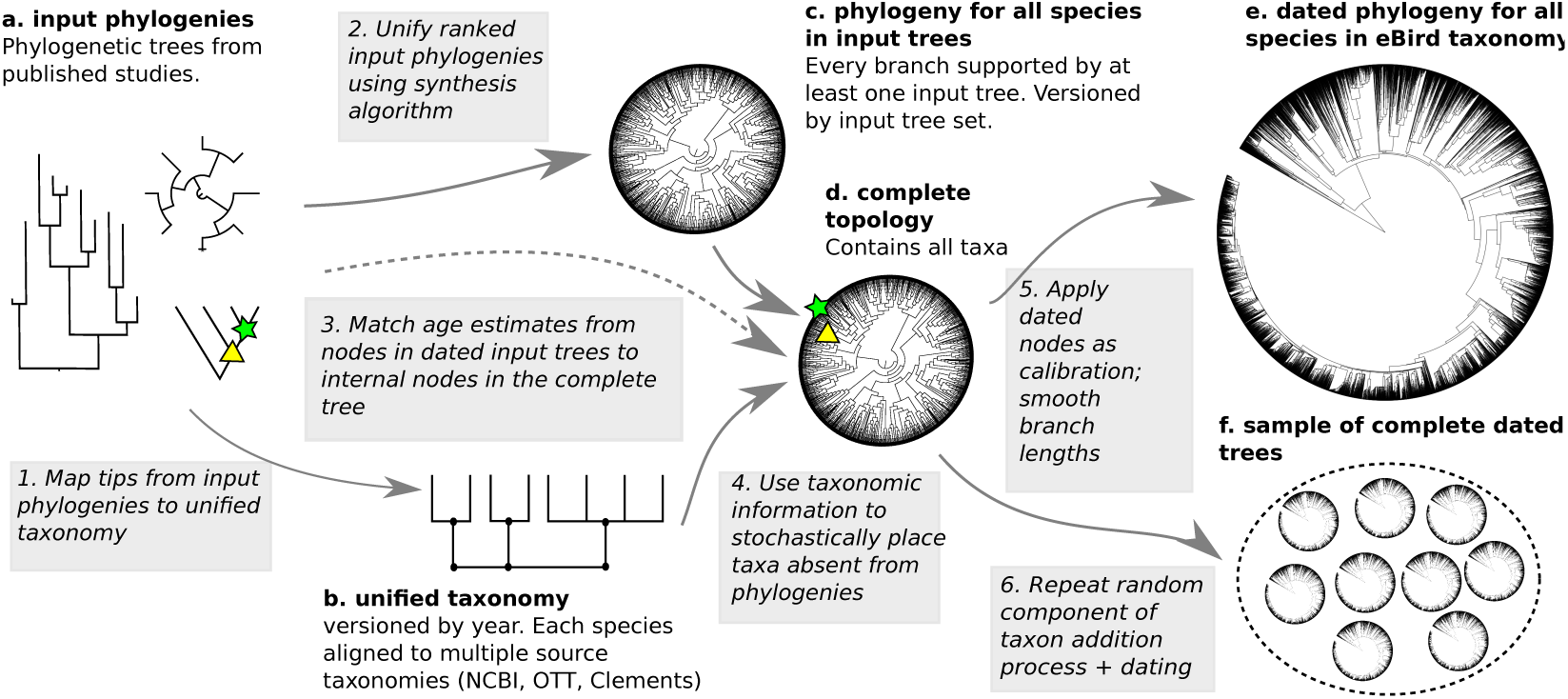
The phylogenetic synthesis workflow. Data products are labelled with letters (a-f) and are shared in the data repositories. Analysis steps are labelled with numbers (1-6) and rely on open source code.

**Figure 2.**
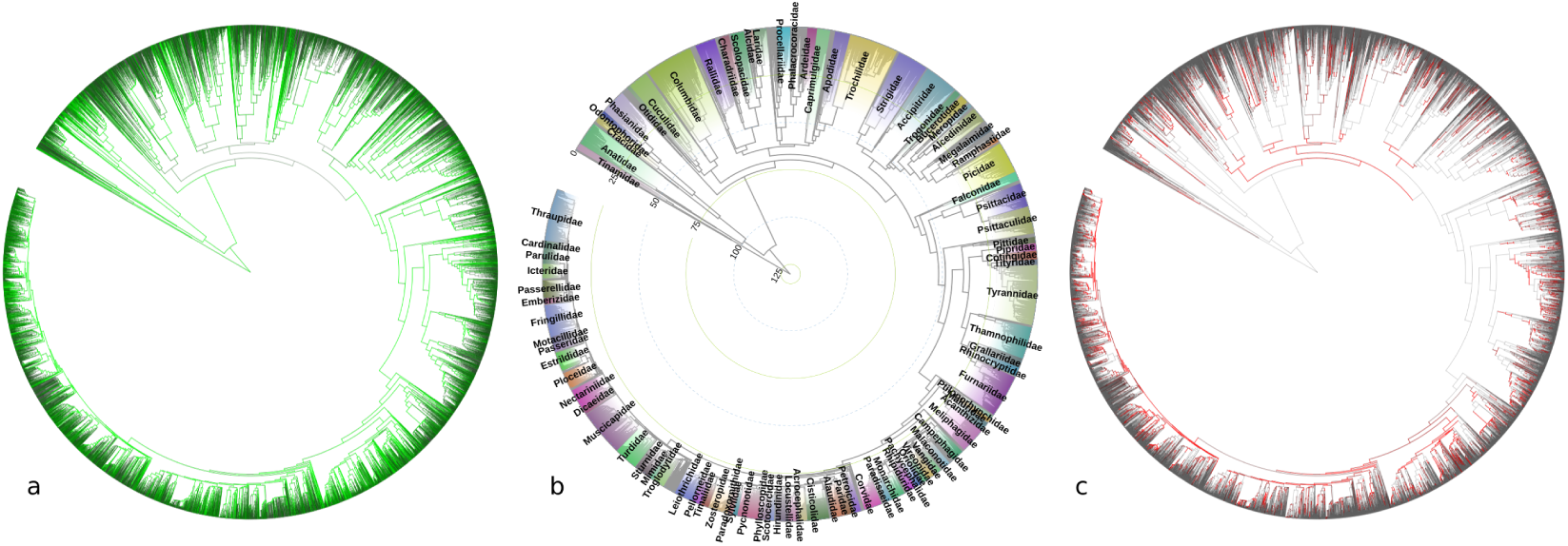
The complete tree labelled with (a) number of trees in support of each branch (grey - no supporting studies; brightest green - 20 or more supporting studies) (b) Bird families with 25 or more members in the tree, and age estimates in millions of years ago (MYA) (c) conflict (grey - no conflicting studies, bright red - up to 15 studies).

**Table 1.** Species count and phylogenetic information across trees aligned to taxonomy versions.

| Clements taxonomy year | Total species count | Species in OTT | Species with phylogenetic information (%) |
| --- | --- | --- | --- |
| 2021 | 10,824 | 10,824 (100%) | 9,239 (85%) |
| 2022 | 10,906 | 10,746 (99%) | 9,207 (84%) |
| 2023 | 11,017 | 10,807 (98%) | 9,189 (83%) |

### Data integration

When data from published papers about evolutionary relationships is shared (which it often isn’t, see [5, 19] ), this data may be in any of a variety of tree file formats. Nearly all tree files require human curation to map tip labels to shared taxonomic identifiers and to ensure correct rooting and metadata collection. We curated 281 individual trees from 262 published studies and included them in the Open Tree phylesystem datastore [20], (Figure 1). These trees comprise a total of 44,117 individual tips. We developed a taxonomic crosswalk table which contains unique identifiers from the Open Tree taxonomy (OTT v3.6) [28] for all 10,824 species in the Clements 2021 taxonomy, and most species in the 2022 (10,746/10,906; 99%) and 2023 (10,765/11,017; 98%) Clements taxonomy [3] (Table 1). We were able to match 35,534 (86%) of the input study tips to species in the Clements taxonomy via identifiers in OTT. Upcoming versions of OTT will include all species from the 2022 and 2023 Clements taxonomies. We used Avibase taxon concepts associated with species in the Clements taxonomy to link identifiers for taxa whose names changed across years [16].

We had phylogenetic information, as defined by a species appearing in at least one input tree, for 9,239 of the 10,824 species in the Clements 2021 taxonomy (2022: 9,207/10,906; 2023 9,189/11,017 (Table 1)). Some input tree labels could not be matched to some versions of the taxonomy, resulting in small differences in the counts of species for which we have phylogenetic information across taxon-years. New phylogenies and corrected name mappings can be contributed to the data store by any user at any time [20].

### Phylogenetic synthesis

All internal nodes in the phylogeny-only tree are supported by at least one input phylogeny (Figure 1c). We publish with the synthesis tree an annotation file tracing the provenance of every branch in the synthetic tree, including what input study and tree supports that branch’s inclusion in the final tree, what trees have branches that align with that branch but do not directly support that branch, and what trees contain relationships that conflict with that branch in the tree [26].

This synthesis tree is updatable as new data become available. We version the tree, and the provenance information for each version includes a static record of the set of inputs and rankings used to generate each tree. The synthesis version described here is tagged as “Aves1.2”, and each subsequent update will receive its own tagged version number. New versions of the tree will be added to the data store available on GitHub and published with a DOI on Zenodo.

The majority of branches in the tree were supported by one to five phylogenies, although some nodes in the tree are supported by up to 24 input phylogenies (Figure 2a). In a testament to the history of change in the estimated phylogenetic relationship of Aves over time, 3,781 branches (around 34%) conflict with at least one study. (Figure 2c). While in most cases of conflict there are one or two input studies that conflict with the resolution at a branch, there are 20 branches with 10 to 15 studies that show an alternate resolution for that relationship. These persistent conflicts may result from different inferences for trees using different data types [25]

To create a complete tree from the phylogenetic synthesis tree, we used a curated taxon addition step. To add taxa that we did not have phylogenetic information for we used the R package ‘addTaxa’ [17] (Figure 1 step 4). Taxa without phylogenetic information are added to the tree using curated taxonomic information files. These taxon information files provide a constrained region of the tree that experts believe the taxon should be placed in, using multiple data sources including Birds of the World [3]. These files are versioned, and readily accessible and updatable. These taxa are then added stochastically based on these constraints. We sampled 100 random taxon addition trees per taxonomy year and, after dating the trees, summarized these sampled trees into a single maximum clade credibility tree. This tree summarizes the relationships into a single complete tree including all taxa. We used this process to generate a complete tree for each of the 2021, 2022, and 2023 versions of the Clements taxonomy.

### Concordance and conflict

This synthesis tree largely agrees with the named relationships as captured by the Clements taxonomy. However, the monophyly of some groups supported by phylogeny conflict with relationships in the Clements taxonomy, although many are been resolved in subsequent revisions, e.g., from Clements 2021 to Clements 2022 and Clements 2023 (Figure 3). While the relationships in the Aves v1.2 phylogenetic synthesis tree remain the same across all 3 taxonomy versions, some names assigned to the tip taxa, and the higher taxonomy associated with these names, change with the taxonomy version. The changes in the number of tips represented in the phylogenetic synthesis are due to taxon name matching changes and pruning the tree to the species level (Table 1). Across years the same taxon may be split into multiple species, or multiple species may be lumped into a single one.

**Figure 3.**
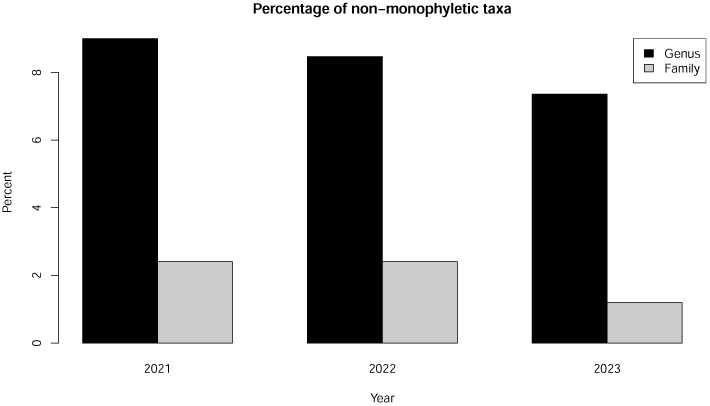
Percentages of taxa, families, and genera, in the Clements taxonomy which are not monophyletic in the to the phylogenetic synthesis tree, across different taxonomy years.

Where taxonomy and phylogeny conflict, some of these disagreements are minor (e.g. non-monophyly of the genus *Bolborhynchus*), while others are more significant. This complete tree captures phylogenetic conflicts with the monophyly of six families, as they were defined in Clements 2021; Muscicapidae, Turdidae, Laniidae, Macrosphenidae, Sarothruridae, and Rallidae. However, in the Clements 2023 taxonomy the monophyly of three of these families was rectified. The taxa which belong to Sarothruridae and Rallidae were updated, and these families are now concordant with the phylogenetic relationships observed. Two species, *Rallina forbesi* and *Rallina mayri*, were taxonomically placed in Rallidae in 2021 and 2022. These taxa rendered Rallidae non-monophyletic with respect to Sarothuridae [9]. In the 2023 Clements taxonomy these species names were updated to *Rallicula forbesi* and *Rallicula mayri* in the family Sarothruridae (Flufftails), restoring the monophyly of both Sarothuridae and Rallidae. Taxonomic updates in 2023 also resolved the non-monophyly of Macrosphenidae; Grauer’s Warbler *Graueria vittata*, the only member of its genus, was reclassified from Macrosphenidae into the family Acrocephalidae (Reed Warblers and Allies) [23], and the genera *Hylia* and *Pholidornis*, which are also monotypic, containing *Hylia prasina* and *Pholidornis rushiae*, were moved to the family Hyliidae. Together these revisions render Macrosphenidae monophyletic in the synthetic tree. Because of evidence that it belongs in Turdidae [8], we used taxonomic information to add the genus Pinarornis to that family instead of Muscicapidae. Reclassifying that species in Clements 2024 will fix the monophyly of both of these families. Laniidae also remains non-monophyletic in the 2023 taxonomy. Based on the 2023 taxonomy, the Crested Jayshrike (*Platylophus galericulatus*) is placed as the sole member of the family Platylophidae. However, per the phylogeny in McCullough et al. 2023 [18] this species is nested within Laniidae.

We made comparisons between our synthetic tree and the Jetz et al. 2012 [12] global bird tree. The Jetz et al. tree [12] has been widely used for large scale bird phylogenetics, and continues to be used as a phylogenetic backbone, even more than a decade later (e.g. [4]). The published tree from Jetz et al. [12] contained 9,993 tips, 6,670 of which were informed by genetic data (66%). We were able to align 9,887 of the 9,993 (99%) tips in the complete tree to Open Tree taxon identifiers, and 9,462 (95%) to taxa in the 2021 Clements taxonomy. In our synthetic tree, we have phylogenetic information placing the relationships of 8,414 (84%) of the taxa in [12]. Many relationships remain the same. Indeed, the tree built using genetic data from [12] is one of the inputs into the synthetic tree, and informs many relationships. However, there have also been updates to our understanding of bird relationships since this tree was published. We include 166 papers on bird phylogeny published since 2012 in this analysis, as well as phylogenetic estimates for many species for which there were no genetic data in [12]. There are 2,377 branches in our tree that capture different relationships than the Jetz et al. tree, and there are phylogenetic relationships for 2,451 taxa in this tree which were not present in [12] or in any other previous single study.

### Dates

We used published date estimates from 119 trees from 89 published studies (full citations are listed in Supplemental Date Citations, and posted in the data repository). These studies provided date estimates for 6,984 (∼ 63%) of the internal nodes in the tree. 1,649 nodes had date information provdied by only one input study, 1,265 nodes had two input date estimates, and 3,964 had between 3 and 10 input date estimates. There were up to 38 date estimates for other nodes. The nodes for which there is no published date information are assigned dates equally spaced between the nearest dated parent and child nodes. While there is high variance among studies [31], and individual dates are approximate, these combined analyses provide a broad view of the timing of divergences across the entire bird tree of life. We generated 100 complete dated trees with topologies stochastically sampled via the taxon addition process, and dated using a random sample from the input node age sets. We summarized these trees into a single topology and branch lengths, with confidence intervals on dates. However, these intervals will underestimate date uncertainty, particularly in regions of the tree with few input date estimates. Importantly, the individual date estimates for each node, and the metadata linking those estimates back to published studies, is published with the data files for the synthetic tree, so users interested in investigating uncertainty across dates can perform their own downstream analyses.

### Custom tree synthesis

The input phylogenies used here are all publicly available via the Open Tree data store [20]. The custom synthesis software is also available both via an API and through a simple browser based interface (https://aves.opentreeoflife.org/v3/tree_of_life/launch_ custom). Any interested user can create their own custom synthesis tree for any set of phylogeneies. This synthesis can be performed on trees already present in the Open Tree data store, or by adding new trees to the data store. Phylogenies can be ranked in any preferred order, and a complete synthesis generated by a user for any set of taxa included in the Open Tree taxonomy. We will update the data repository and share new versions of the Aves synthetic tree, at least annually and potentially more often.

### Data attribution

A key premise of our project is the dynamic but reproducible nature of the output. Empowering other researchers to contribute and, critically, be recognized for their own work, is a central goal of the project. Because the provenance of all relationships in the phylogeny are carefully tracked, we are able to readily quantify the proportion of nodes in the final tree that are informed by a given study (Table S1). This remains true for subtrees pruned to represent smaller sets of taxa such as those in a regional study, and we provide easy-to-use code for how to derive the input trees that contribute to the relationships for trees of arbitrary sets of taxa from the larger tree. While we firmly believe that users of our phylogeny should cite all the data sources that go into the phylogeny they end up using for their own study, we recognize that most journals have a limit on the number of citations in an article. Here we have cited all 262 inputs in the supplemental material. The proportions of nodes supported by a constituent study could help users to identify the most critical studies. We hope that in general this project adds to a growing understanding of the need to adequately track the impact of datasets themselves, which may require changes to journal policy. An important corollary is that, although most journals require data deposition, which in some cases explicitly and in most cases implicitly, covers phylogenetic estimates, many authors still do not publish the actual tree files they generate and use for downstream analyses [5, 19] Many phylogenies are available only as images in PDFs. This lack of accessible data precludes others from reproducing results, and is likewise something we hope that journals address. The data access and attribution pieces go hand in hand, and by building the tools to track these linkages, we aspire to move the needle on these important aspects of the modern scientific process. The Open Tree project provides not only a stable and usable data store, it incentivises data curation and sharing by offering tools such as custom synthesis and visualizations for consensus and conflict between trees, and between trees and taxonomy [20].

### Interoperability

To demonstrate the power of data linkages between this project and others, such as eBird [30] –the largest community science project in the world–we downloaded all eBird records, filtered to those we could confidently assign to a precise location, and grouped these into hexagonal grid cells at the global scale. We then calculated the mean pairwise phylogenetic distance among the species in each grid cell, which had the effect of graphically and quantitatively highlighting a number of known but poorly understood global gradients in evolutionary relationships among co-occurring species (Figure 4). For example, the Andes, particularly at high elevations, are characterized by a phylogenetically clustered set of species when compared to lowland Amazon. The pattern is even starker in the Himalayas, where over a relatively short geographic distance the pattern goes from distant relatives co-occurring with one another, to increasingly closely related species co-occurring at higher elevations and into the Tibetan rain shadow [24]. Inputs to eBird data vary and are higher in densely populated areas. Data is limited in areas, almost totally absent from some others, and very dense in others. For example, the linear tracks across the oceans reflect cruise ship routes, and the apparent deep divergences in the UK likely reflect concerted birdwatching effort there uncovering a bevy of odd vagrant species. Nonetheless the deep evolutionary divergences between species in Madagascar, lower elevations in Papua New Guinea, Southern South American, and Southeast Asia are also visible, and these divergences contrast with the profound phylogenetic clustering of arid and high latitude communities in places such as western Australia [22], the Sahara, and the Southern Ocean. Linking patterns such as these with trait data such as those from AVONET [32] will provide insight into the processes structuring global avian diversity, and such approaches will be greatly facilitated by the interoperability of our project with other data resources like those mentioned here.

**Figure 4.**
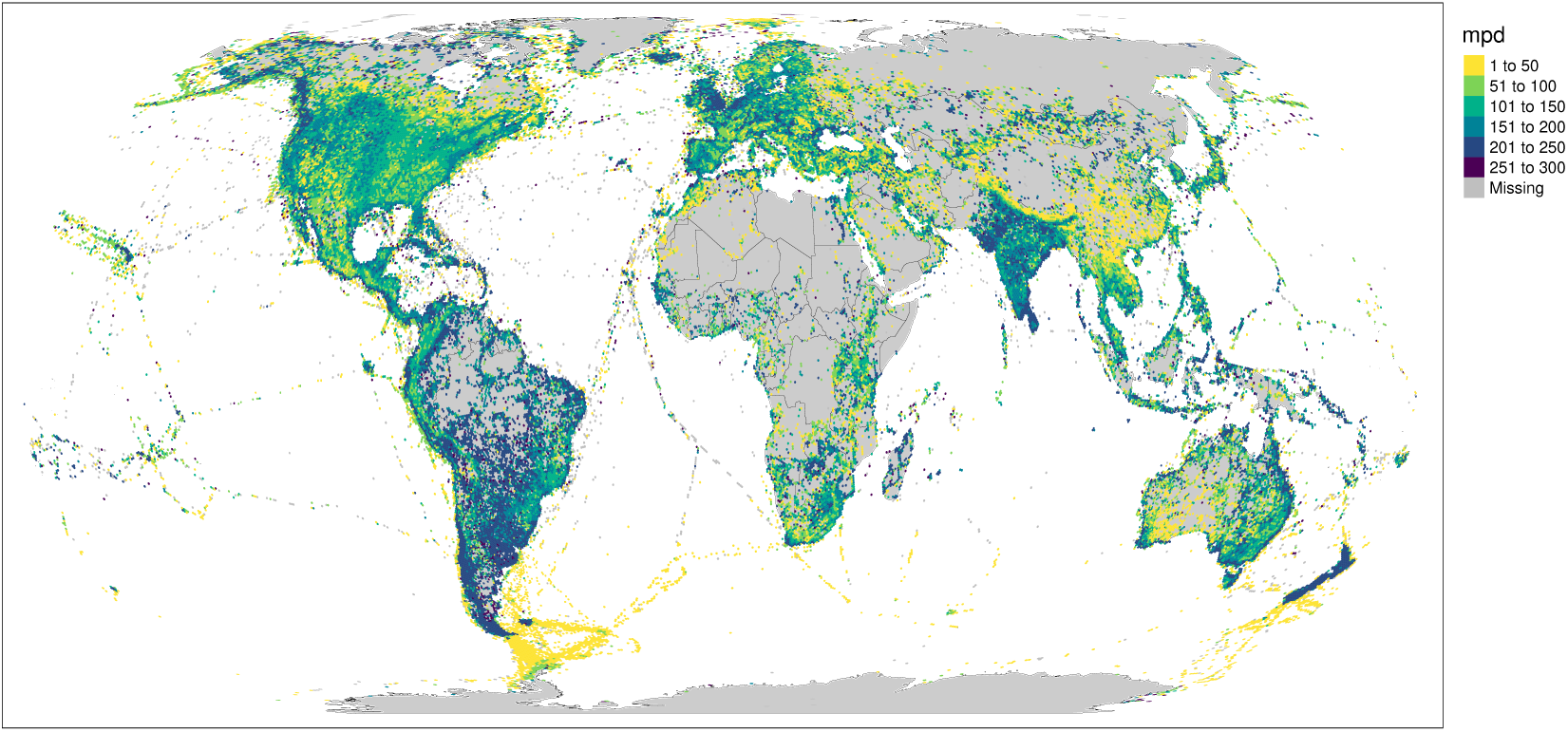
Global phylogenetic diversity of bird records across the globe. Co-occurring bird species tend to be more distantly related in the tropics and in what were once elements of the ancient Gondwanan continent, including Madagascar, New Zealand, South America and India. Grid cells on the map are colored according to global mean pairwise phylogenetic distance among species reported to eBird.

## Discussion

Intrinsically, this synthesis tree approach builds on prior work. We extend and expand the synthesis trees built using earlier versions of the Open Tree synthesis software [1]. The efficiency of this software has been improved, allowing for rapid synthesis of even very large trees [27].

Taxonomy is a critical axis for interoperability between projects. In the past, discordance between the taxon names and concepts captured in the Open Tree and the Clements taxonomies has limited ornithological community engagement with Open Tree resources. The automated algorithms required to create a tree of life at the scale of all the world’s organisms result in an avian phylogeny with more taxonomic inconsistencies and less resolution than ornithologists expected. Unifying taxonomic concepts across different resources and through time is non-trivial, and often requires in-depth knowledge of the literature and the history of research on a given taxon. While some automatic name matching is possible, automated systems can also easily lead to errors. Here, via a curated process, we strike a balance between linking data both to large resources such as NCBI and GBIF, and to taxon concepts in the most up-to-date and accurate bird taxonomic resources [16]. This project fills a key gap in current biodiversity informatics - cross-linking names, and unique identifiers associated with those names, at scale. Mapping taxonomic concepts and names across resources in a public, accessible, and reusable database is essential for leveraging the open data resources being generated by the biological community [11].

The tips of the phylogenetic synthesis tree are in principle at the species level, but different taxonomies vary slightly in whether some taxa defined as species or subspecies. In some cases the definitions for a specific taxon concept also change across years of the Clements taxonomy. Each of the tips in our tree are mapped to identifiers and labels in the Open Tree taxonomy, and in three different versions of Clements taxonomy. This process is facilitated by the use of Avibase taxon concepts [16]. In principle, it would be possible to initially curate trees using these taxon concepts, and this would further facilitate taxonomic updates and minimize attrition of matched taxa over time, but that approach is not currently technically possible.

Our approach also identifies pathways that can be built to link the phylogenetic estimates to the vouchered tissues, physical specimens, and sequence archive records from which they are derived. This interoperability will improve the bioinformatic connections to collections and community science data.

Many birdwatchers keep careful track of which birds they have and have not seen. eBird makes this process fun and easy, and over 800,000 participants have taken advantage of the tool, contributing valuable citizen science data in the process. This tree directly maps to the species lists used in eBird, and provides an evolutionary framework for analyzing those data (Fig. 4). These links between resources create an opportunity to feed evolutionary information back to birders, capturing the phylogenetic diversity of their observations.

The synthesis approach also affords the opportunity to explore concordance and conflict across estimates of bird relationships through time. While most relationships have remained stable, others are highly contentious [25]. In our consensus tree we use a very simple ranking system - relationships in more recent trees are preferred over those in older trees (with the exception of two very large super matrix trees with relatively high levels of missing data, Burleigh et al. 2015 [2] and Jetz et al. 2012 [12] which were ranked last.) This metric captures the ideal that we are getting closer to accurate estimates of relationships through time.

While few groups are as well studied and data-rich as birds, the approaches described here can be applied to unifying taxonomic and phylogenetic information for any group. Our synthesis provides an evolutionary framework to harness trait data, phylogenetic inferences, collections data, observation records, and taxonomic concepts, in shareable, reproducible, interoperable datastores.

## Materials and Methods

### Data integration

We selected and curated 281 trees from 262 published studies containing bird phylogenies published from 1990 through 2024. All of these trees and their associated metadata are now available in the Open Tree data store, phylesystem [20]. The full set of citations are posted with the data at https://github.com/McTavishLab/AvesData and are in the supplemental materials.

We gathered phylogenetic trees associated with publications from online data stores, and by contacting authors directly. The metadata for each tree was curated to contain standard phylogenetic metadata comprising ’Minimum information about a Phylogenetic Analysis’ (MIAPA) data [15]. These data and metadata are stored in an open data store, in an JSON format translation of NeXML [33]. The data store is mirrored to GitHub, and available open access. The datastore is editable by curators, and all edits and updates are tracked as git commits.

We mapped tips in trees to identifiers in the Open Tree taxonomy (OTT). The Clements taxonomy is used by the Cornell Lab of Ornithology, which manages eBird, among other data resources. To link taxon names across Clements and OTT, we created a taxonomic translation table which maps every species level taxon in the 2021 Clements taxonomy to a unique identifier in OTT. Via OTT, our translation table links identifiers from the Clements taxonomy to NCBI, GBIF, Catalog of Life and other online taxonomic resources, and is used by the Open Tree project [28]. Taxa can be added to OTT via an amendment system. The taxonomic crosswalk is shared with the data resources for this project on GitHub. To match taxon names we used the Open Tree bulk taxon name resolution system, which searches on canonical names and on synonyms stored in OTT https://tree.opentreeoflife.org/ curator/tnrs/. Synonyms in OTT are imported from the component taxonomic resources, primarily NCBI and GBIF. Additional matches were made by hand by the authors of this manuscript. Whether matches were made based on ‘canonical name’, ‘synonym’ or ‘hand match’ is tracked on the crosswalk table. We added species that were in the Clements taxonomy, but not present in the OTT to OTT via the taxon amendment process. These taxa are available in OTT v 3.5. For each of the input trees, we mapped tips to unique OTT identifiers.

We added some taxonomic constraint trees to reinforce monophyly of higher level taxa from Clements where there were no direct phylogenetic conflicts with the monophyly of those groups. The Open Tree synthesis algorithm preserves monophyly of taxa where there is no phylogeny directly contesting that relationship, but due to some differences in the membership of groups in the higher taxonomy of the Open Tree taxonomy as compared to the Clements taxonomy, we added constraints capturing those taxonomic groups to ensure those taxa were present in the synthesis tree. All trees used in this analysis are publicly stored in phylesystem in the collection ‘snacktavish/Aves’.

### Phylogenetic synthesis

We ranked the input trees by publication year, with more recent trees ranked higher, with the exception of two large supertrees built from sparse matrices, Burleigh et al. 2015 [2] and Jetz et al. 2012 [12]. We ranked these trees below the other published phylogenies.

We performed custom synthesis using the Open Tree synthesis software. To create a custom synthesis tree, we combine input trees with the taxonomic backbone constructed from a single reference taxonomy using the propiquinty and otcetera software developed for the Open Tree of Life project and described in depth in [26]. The taxonomy provides a placeholder for taxa that we have no phylogenetic information on. Taxon names on input trees are standardized to the same reference taxonomy that was used to construct the taxonomic backbone. This process uses a greedy algorithm that adds branches from input trees in rank order, from the mostly highly ranked tree, to the lowest ranked tree. To make supertree construction tractable, we decompose the overall task into sub-problems, based on monophyletic taxa [26]. We look for named groups in the pruned taxonomic backbone that are “uncontested” by any single phylogenetic tree. This allows for subdividing the tree in smaller sub-problems that will be easier to solve using phylogenetic evidence. To solve the sub-problems, there are two steps. First, evidence for each sub-problem is identified among exemplified phylogenetic trees and the pruned taxonomic backbone. This is done by pruning away all tips that do not belong to the sub-problem from the exemplified phylogenetic trees and the pruned taxonomic backbone. Then, the ranking of phylogenetic trees is used to sequentially gather evidence to resolve relationships in the sub-problem. Taxonomic evidence is always ranked last, below any published phylogenetic estimates. Concordance and conflict between each input tree and the resulting estimate is stored, and output as synthetic tree metadata. Recent tooling developments in otcetera [27] have sped up this synthesis process, so that updated synthesis trees for all of Aves (around 11,000 taxa) can be estimated in a few minutes. Every branch in the synthetic tree is annotated with the node in an input study or the taxonomic relationship that supports that branch. These links from input data to each relationship in the synthetic tree make the route from input data to inference clear, and provide a direct path to correcting incorrect inferences.

We created versions of the synthetic tree aligned to the 2021, 2022 and 2023 updates to the Clements taxonomy in our GitHub repository https://github.com/McTavishLab/ AvesData.

### Adding taxa without phylogenetic information

We prune the synthetic tree to only taxa that are included in input phylogeneics, and places the remaining taxa using a curated taxon addition process (Figure 1 step 4). This process uses modified version of the addTaxa algorithm ( [17];https://github.com/eliotmiller/ addTaxa), which we describe in brief here. First, missing taxa are identified. ’Missing taxa’ are any species in the Clements taxonomy for a given year which are not included in the phylogenetic synthesis tree. Second, taxon addition statements are generated for each missing taxon. Each statement takes the form of a family, genus, or species groups (i.e. a clade, as identified by the most recent common ancestor of two or more species passed to the algorithm). Multiple addition statements can be provided, and the algorithm will search for the most specific placement possible. For example, in Figure S1, both members of the genus Poliocephalus are missing from the grebe family tree. The first Poliocephalus is placed at the family-level, while the second is added as sister to the first. Importantly, these taxon addition statements can also contain exclusion statements. Again, using the example in Figure S1, we specify that Poliocephalus should not break the monophyly of any of the other grebe genera. Third, we implement these taxon statements by randomizing the order missing species are added in, then iterating the process multiple times to generate a cloud of 100 possible, taxonomically complete trees (Figure 1 step 6). By applying this post-processing step, we are able to generate from each phylogenetic synthesis tree a distribution of complete, bifurcating, phylogenies, with branch lengths proportional to time.

### Estimating dates

There is no branch length information in the custom synthesis tree because the branches are combined estimates from phylogenies generated using a variety of data types and inference types, there is very little information about relative node ages for which there are no secondary calibrations. We applied date estimates to the complete trees using Chronosynth https://github.com/OpenTreeOfLife/chronosynth. The Chronosynth approach is expanded from the concepts used in Datelife [31], an R package to estimate dated trees using chronograms in the Open Tree data store [20]. Chronosynth is written in python and relies on python-opentree as a wrapper for Open Tree API calls [21]. Chronosynth summarizes the date information available in the Open Tree datastore phylesystem [20] by mapping the date estimates from nodes in dated input trees to the nodes in the custom synthetic tree which align with those dated nodes. We map aligning nodes using the tree to tree conflict functionality in otcetera [26]. Each internal node in the custom synthetic tree contains a list of source tree nodes that support the synthetic tree node. Imagine pruning the synthetic tree down to have the same terminal set of species as an phylesystem tree and suppressing all internal nodes in the induced tree which have only one child node. If one of the internal nodes, *X*, of this induced tree is the ancestor of the exact same set of species as some internal node, *Y* , in the phylesystem source tree, then node *X* maps to node *Y* . If the parent of node *X* is retained in the induced tree (not removed due to having only one child), then we say that *X* is supported by *Y* . See the discussion of the “supported by” annotation in [26] for further details. This alignment step means that when the topology of the custom synthesis tree differs from that of the dated input tree, not all node dates from the dated input may be used in date inference on the complete tree. Dates which are mapped from the input trees to the synthetic tree then are used as node calibrations. The confidence intervals from input trees are not currently incorporated into Chronosynth estimates. Once nodes are mapped from all of the dated input trees, each node in the custom synth tree may have zero, one, or many dates associated with it. Where multiple dates are available for a node that can be summarized using the a mean, or randomly re-sampled across iterations of summarization.

To sample across some of the uncertainty in dates we estimated smoothed complete trees for each of our 100 sampled taxon addition trees, using a random sample from the node dates for each node. We used a root age for aves of 130 million years ago (MYA) for our smoothed trees. This root age estimate aligns with that in Wu et al 2024 [36], and is slightly older than earlier estimates of 60-118 MYA across earlier analyses [14]. We summarized the node dates using bladj [34]. We share these 100 dated trees in the data repository. We then summarized this set of trees using TreeAnnotator [6] to get a summary topology, and confidence intervals on some dated nodes, which is also shared in the data repository. Trees dated using Chronosynth should be considered rough date estimates, that summarize the existing information about node ages across many taxa and studies, rather than providing novel node age information.

### Taxonomic updates

We published our data package with with the synthesis tree mapped to the Open Tree taxonomy and to the species in Clements taxonomy versions 2021, 2022 and 2023. While most of the taxonomy has remained stable, there have been a few hundred taxon additions and deletions over these three years. We will continue to update and publish new versions of the synthetic tree estimates, as new bird phylogenies are published and added to the data store. In addition, we will share and publish versions of the tree matched to new versions of the Clements taxonomy.

## Acknowledgements

EJM acknowledges NSF grant#1759846. MTH acknowledges NSF grant#1759838. ETM acknowledges support from Schmidt Futures via an Eric and Wendy Schmidt AI in Science Postdoctoral Fellowship to Cornell University.

## Supplemental Materials

**Figure S1:**
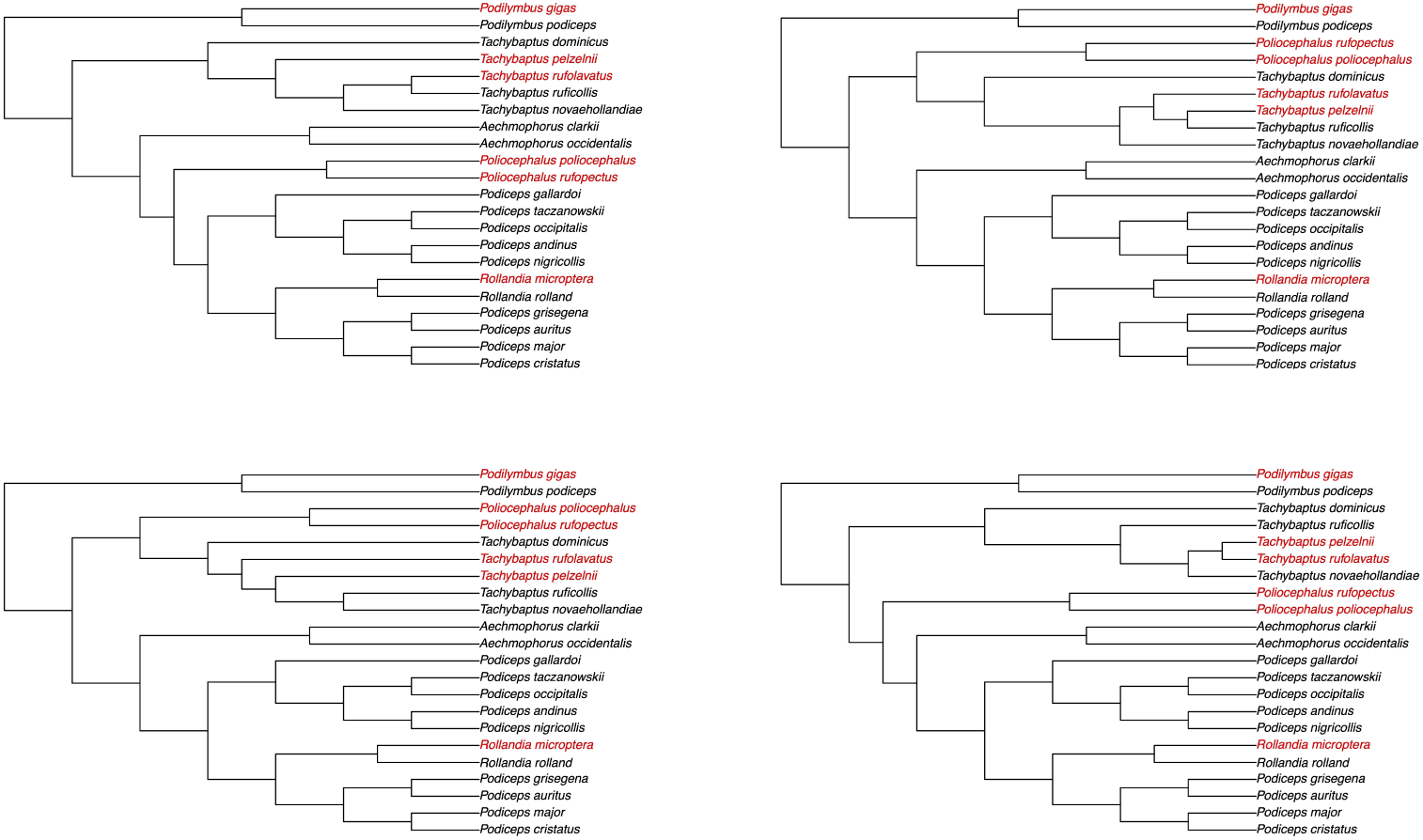
Example trees demonstrating randomization within taxonomic constraints during the taxonomic addition process. Taxonomic constraints were developed from information in Birds of the World

**Table S1.**
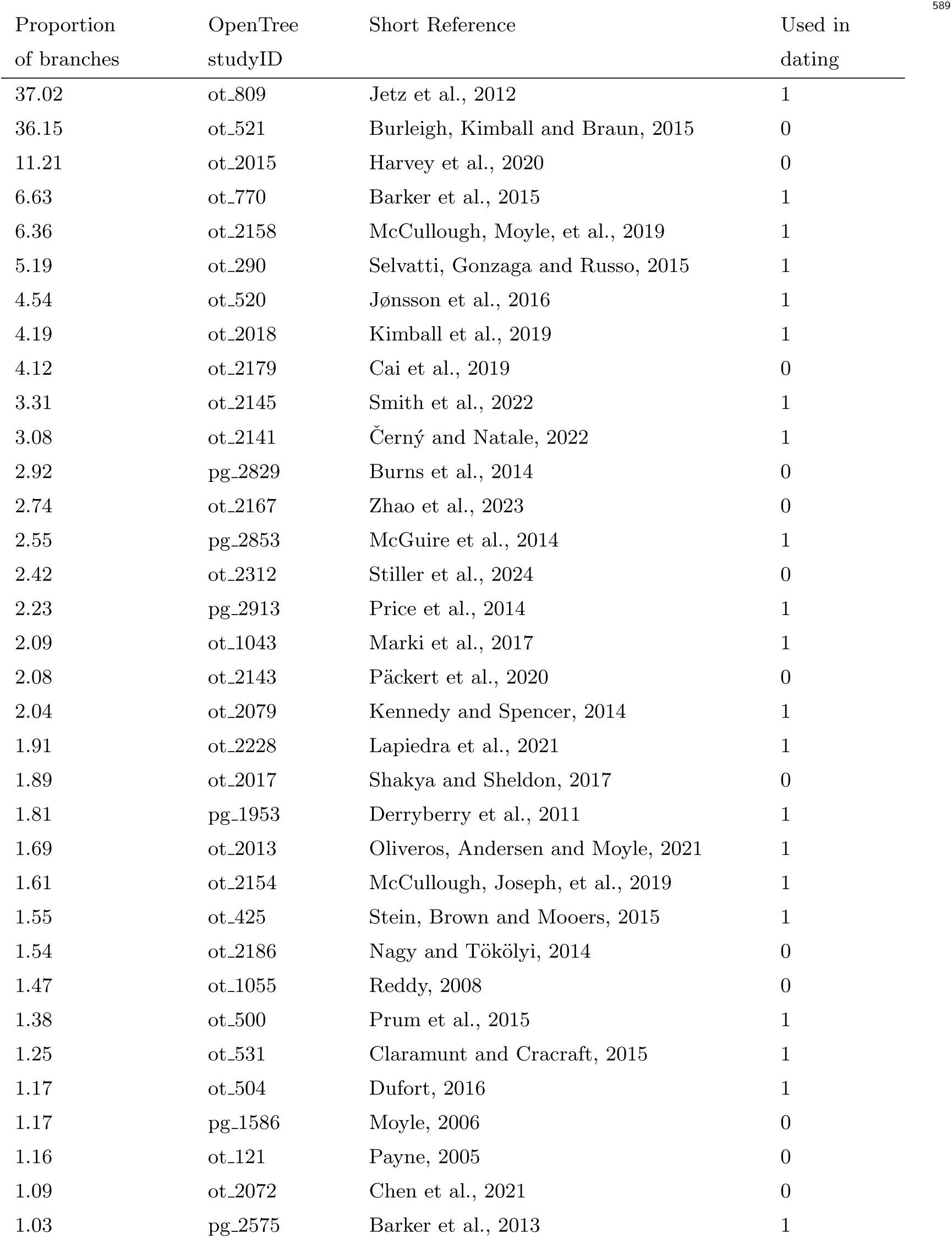
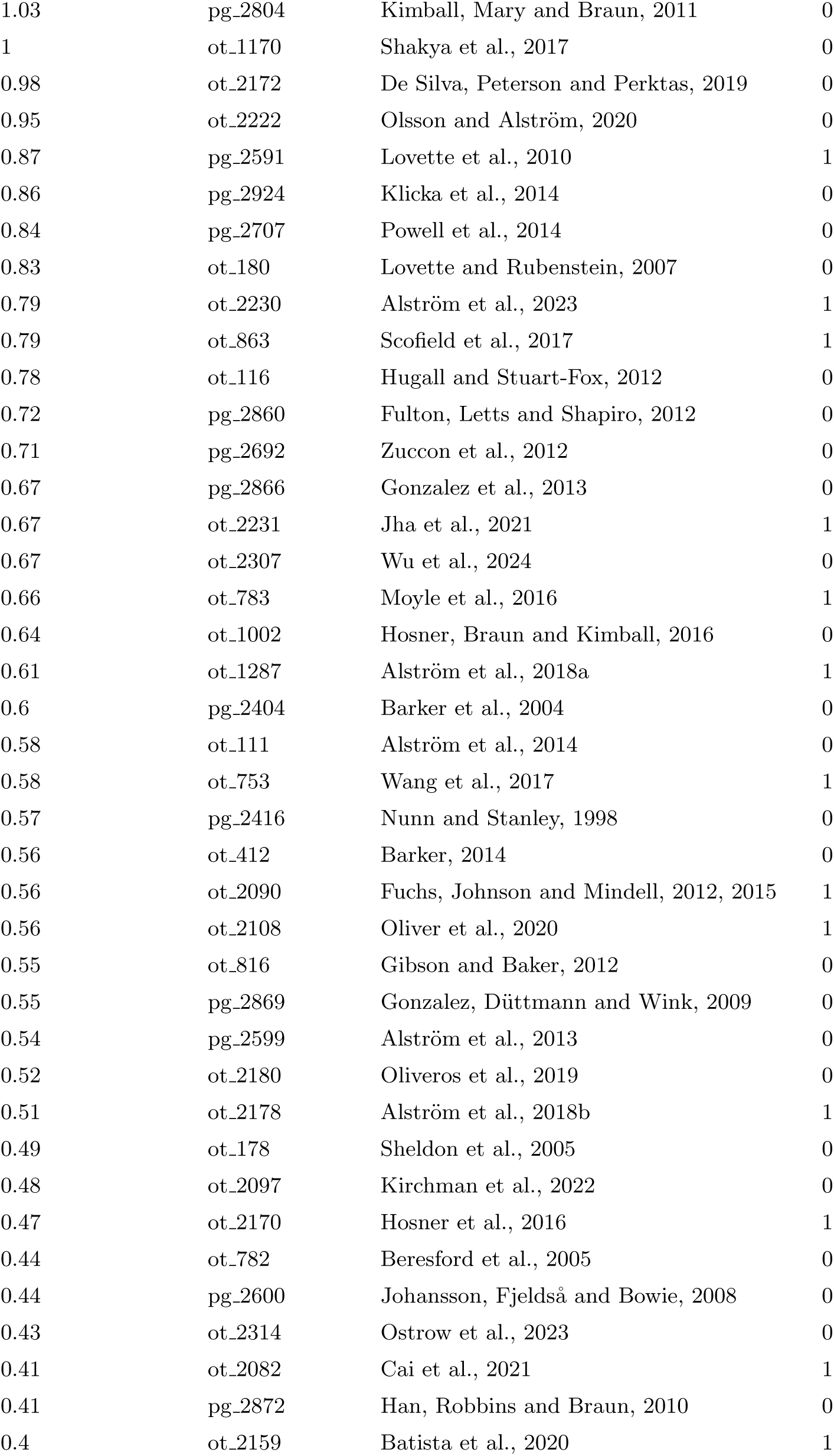
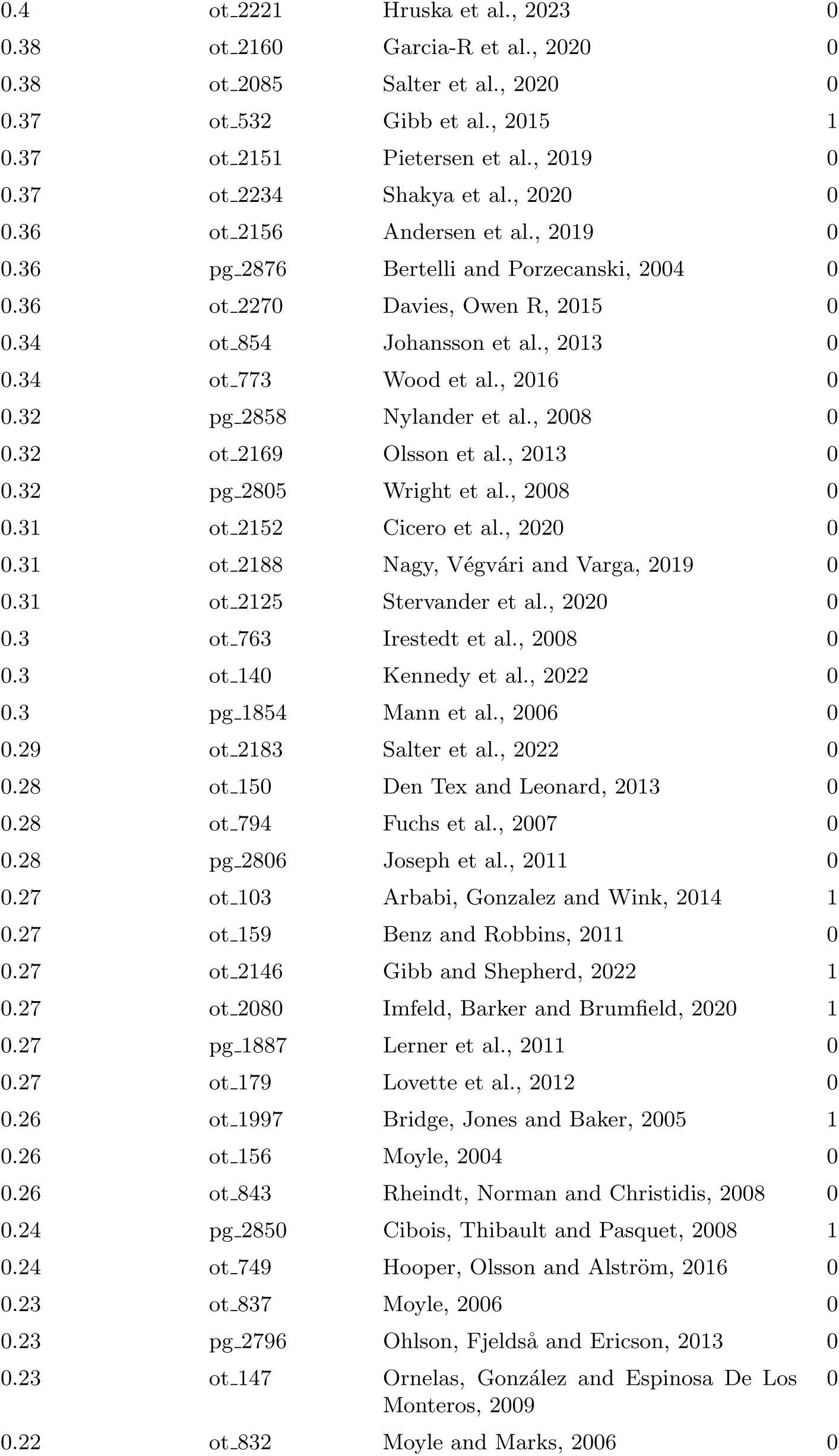
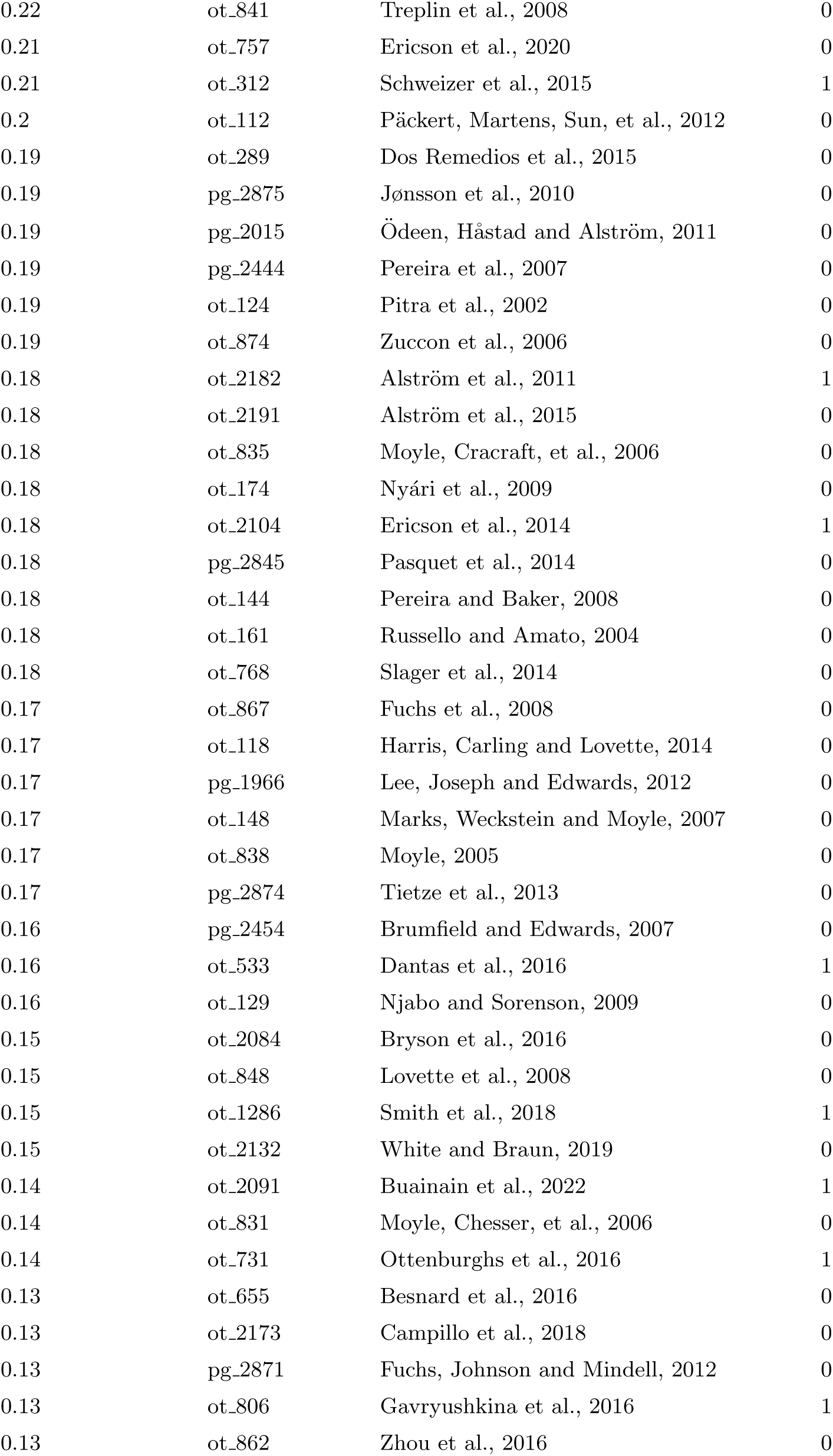
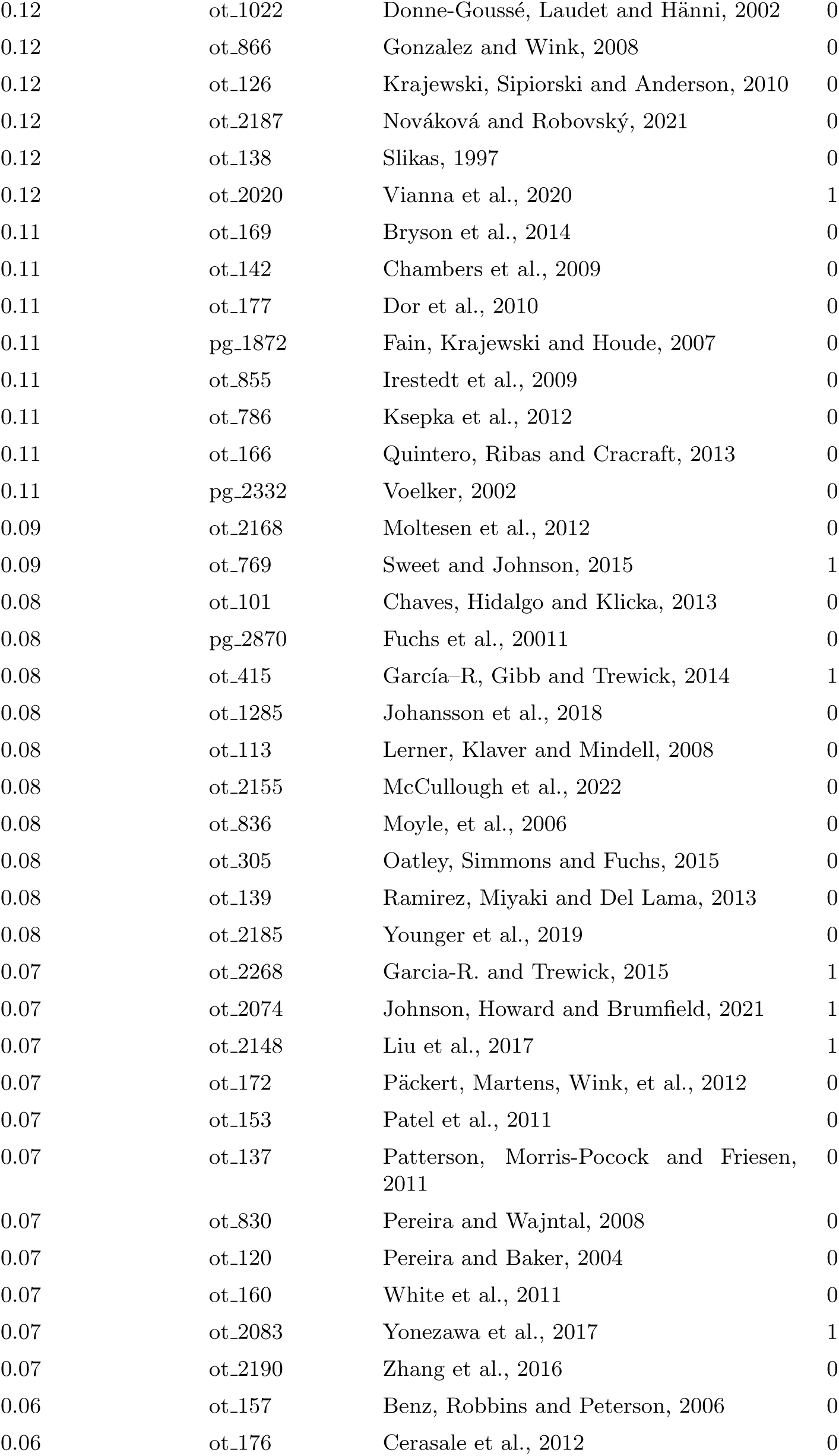
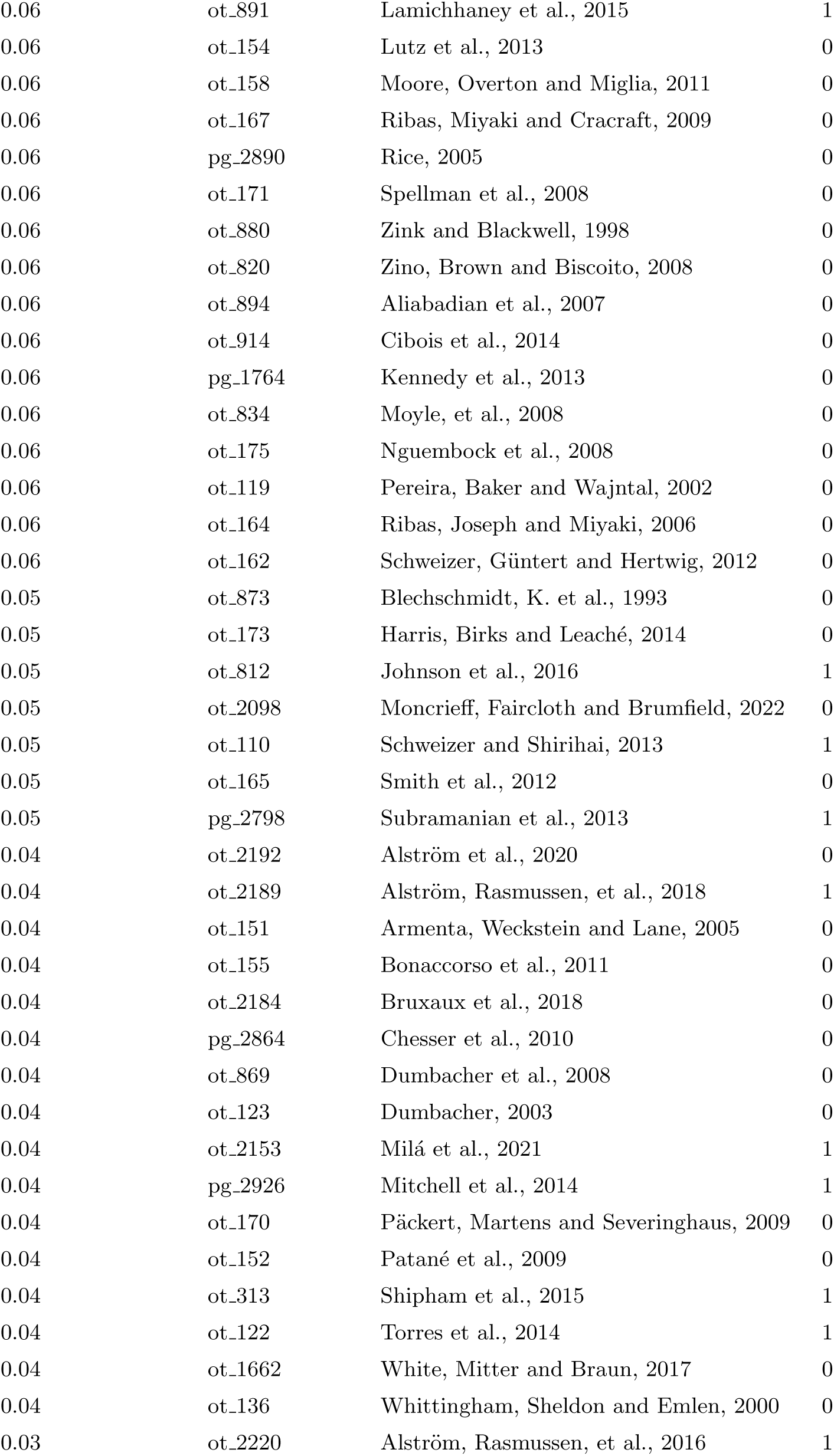
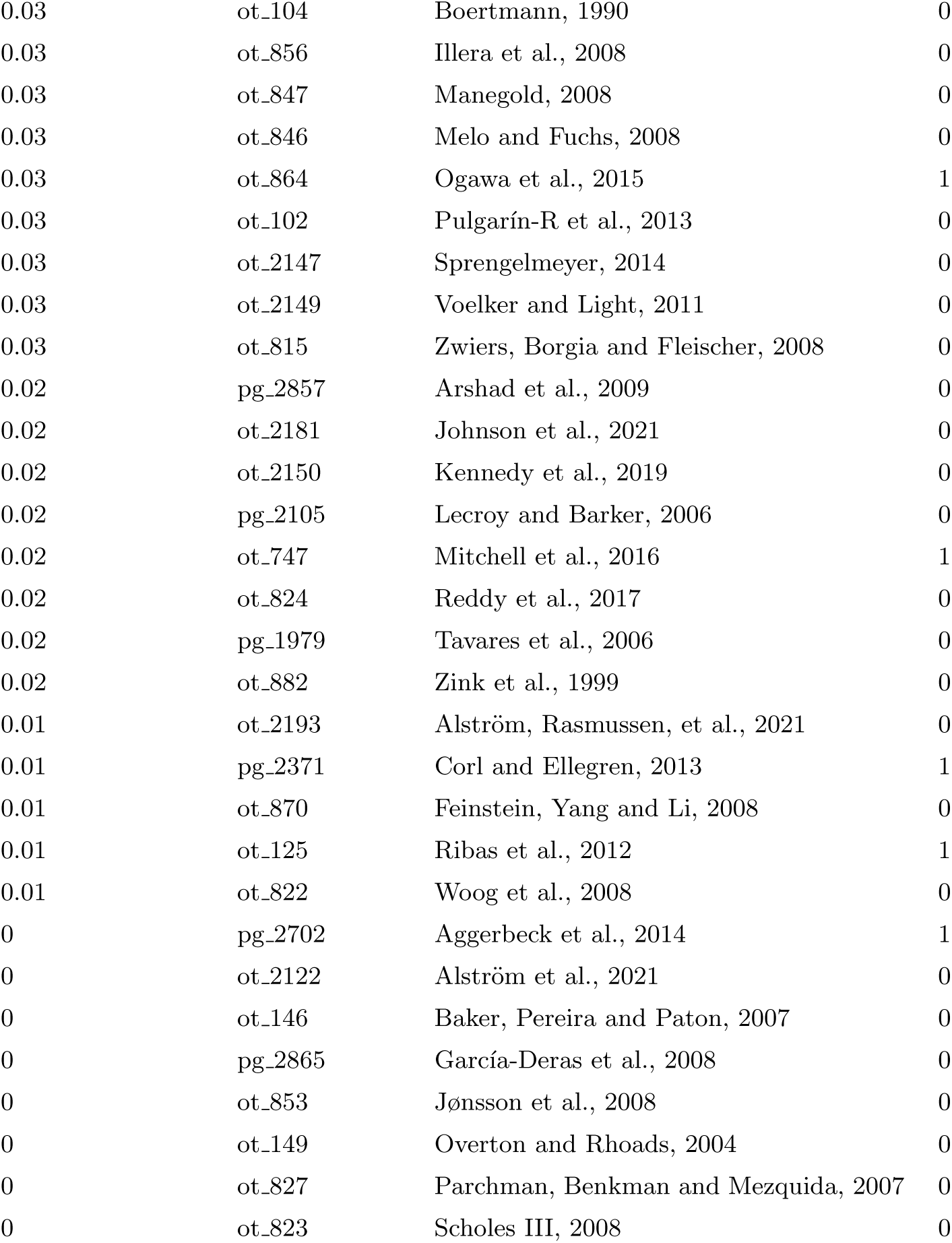
Table showing the proportion of branches in the complete tree supported by each input phylogeny. Column ’used in dating’ shows if dates from that study were included in sampling of internal node date estimates.

